# A candidate gene cluster for the bioactive natural product gyrophoric acid in lichen-forming fungi

**DOI:** 10.1101/2022.01.14.475839

**Authors:** Garima Singh, Anjuli Calchera, Dominik Merges, Jürgen Otte, Imke Schmitt, Francesco Dal Grande

## Abstract

Natural products of lichen-forming fungi are structurally diverse and have a variety of medicinal properties. Despite this, they a have limited implementation in industry, because the corresponding genes remain unknown for most of the natural products. Here we implement a long-read sequencing and bioinformatic approach to identify the biosynthetic gene cluster of the bioactive natural product gyrophoric acid (GA). Using 15 high-quality genomes representing nine GA-producing species of the lichen-forming fungal genus *Umbilicaria*, we identify the most likely GA cluster and investigate cluster gene organization and composition across the nine species. Our results show that GA clusters are promiscuous within *Umbilicaria*, with only three genes that are conserved across species, including the PKS gene. In addition, our results suggest that the same cluster codes for different but structurally similar NPs, i.e., GA, umbilicaric acid and hiascic acid, bringing new evidence that lichen metabolite diversity is also generated through regulatory mechanisms at the molecular level. Ours is the first study to identify the most likely GA cluster, and thus provides essential information to open new avenues for biotechnological approaches to producing and modifying GA and similar lichen-derived compounds. We show that bioinformatics approaches are useful in linking genes and potentially associated natural products. Genome analyses help unlocking the pharmaceutical potential of organisms such as lichens, which are biosynthetically diverse but slow growing, and difficult to cultivate due to their symbiotic nature.

**Importance:** The implementation of natural products in the pharmaceutical industry relies on the possibility of modifying the natural product (NP) pathway to optimize yields and pharmacological effects. Characterization of genes and pathways underlying natural product biosynthesis is a major bottleneck for the use of natural products in the pharmaceutical industry. Genome mining is a promising and relatively cost- and time-effective approach to exploit unexplored NP resources for drug discovery. In this study, we identify the most likely gene cluster for the lichen-forming fungal depside gyrophoric acid in nine *Umbilicaria* species. This compound shows cytotoxic and antiproliferative properties against several cancer cell lines, and is also a broad-spectrum antimicrobial agent. We identify the putative GA cluster from nine *Umbilicaria* species. This information paves the way for generating GA analogs with modified properties by selective activation/deactivation of genes.

## Introduction

Natural products (NPs) and their derivatives/analogs constitute about 70% of modern medicines (1, 2). NPs alone, however, i.e., unmodified molecules as produced by organisms themselves in nature, constitute only a small portion of this. The vast majority, about 60-65%, are derivatives and analogs of naturally-occurring substances, synthesized through biotechnology or synthetic approaches (2, 3). The use of NPs in the pharmaceutical industry relies on the ability to modify NP pathways in order to optimize yields and pharmacological effects. Culture-dependent approaches to identifying/producing NPs are labor-intensive and time-consuming, and not successful for every organism (4, 5). As a result, the biosynthetic potential of many biosynthetically prolific organisms remains untapped. Information on the genetic background and mechanisms of NP synthesis may thus contribute to fast-track NP-based drug discovery (2, 6).

Lichens, symbiotic organisms composed of fungal and photosynthetic partners (green algae or cyanobacteria, or both at the same time) (7–9), are a treasure chest of NPs (10–12). So far, about 1,000 NPs with great structural and functional diversity have been reported from lichen-forming fungi (LFF), and about 300-400 have been screened for bioactivity (11). However, the genetic background of more than 97% of lichen NPs is unknown (13–16). Lichen compounds have great pharmacological potential, encompassing antimicrobial, antiproliferative, cytotoxic and antioxidant properties (11, 17–20). However, there are various major bottlenecks for using lichen NPs in the pharmaceutical industry, including low yield in nature, slow growth, tedious isolation/culturing methods, and a lack of understanding of their genetic background. Targeted genome mining approaches integrate the latest DNA sequencing technologies with computational advancements and large, publicly-available databases of pre-identified BGCs to characterize genes coding for NPs (1, 21, 22). This approach combines genome mining with the expected genetics of the NP to narrow down the candidate biosynthetic genes.

In-silico approaches for linking natural products with their respective biosynthetic gene clusters (BGCs) – genomic clusters of biosynthetic-related genes typically found in fungi (23–25) – are becoming more common in LFF due to the increased availability of genomic resources and databases (1, 21, 22), improvement of detection software and genome mining tools, stabilizing PKS phylogenies, and information gained from recent successes in the heterologous expression of *PKSs* from LFF (13, 14). For instance, the clade “Group I, PKS 16” from Kim et al (13) is associated with the biosynthesis of orsellinic acid derivatives (orcinol depsides and depsidones) such as lecanoric acid (14), grayanic acid (15), physodic acid and olivetoric acid (16), whereas the clade “Group IX, PKS23” from Kim et al (13) is associated with the biosynthesis of methylated orsellinic acid derivatives (β-orcinol depsides and depsidones) such as atranorin. The cluster linked to usnic acid biosynthesis is also fairly well understood (26, 27) and corresponds to “Group VI, PKS8” from Kim et al (11).

Here, we combine high-throughput long-read sequencing with a comparative genomics approach to identify the putative cluster(s) linked to the synthesis of gyrophoric acid (GA). GA is an NP produced by several LFF species, with a broad spectrum of bioactivity (pharmacological properties such as anticancer and antimicrobial activity and industrially-relevant properties such as usage as dyes (19, 28–30)), for which the molecular mechanism and genetics of synthesis remain unknown. Identification of the GA gene cluster would facilitate its production via biotechnology to optimize the yield as well as to generate GA analogs with the desired pharmaceutical effect. For this study, we chose nine species of GA producers belonging to the lichen-forming fungal genus *Umbilicaria* (Table 1). GA is the most characteristic compound of this genus, and is found at high concentrations in all the chosen species (28, 31–33). It is a depside containing three orsellinic acid-type rings joined together by ester bonds (Fig. 1). Apart from GA, several other structurally-related depsides such as umbilicaric acid, lecanoric acid and hiascic acid (Fig. 1) have also been reported from *Umbilicaria*, but these usually constitute a minor fraction (<10%) of the total NPs detected via HPLC (Fig. 1).

**Table 1.** Genome quality and annotation statistics.

| Table 1 Genome quality and annotation statistics |  |  |  |  |  |  |  |  |
| --- | --- | --- | --- | --- | --- | --- | --- | --- |
| Species | TBG number | ccs HiFi yield | # scaffolds | N50 | Completeness | assembly size (Mb) | # Genes | # Proteins |
| *Umbilicaria freyi* | TBG_2329 | 47.39% | 113 | 2575640 | 95.7 | 53.4 | 10,156 | 10,065 |
| *Umbilicaria freyi* | TBG_2330 | 46.41% | 54 | 2043163 | 85.9 | 50 | 8,848 | 8,773 |
| *Umbilicaria deusta* | TBG_2334 | 47.86% | 47 | 1669916 | 97.6 | 41.6 | 8,949 | 8,857 |
| *Umbilicaria deusta* | TBG_2335 | 43.54% | 42 | 1865302 | 90.2 | 37.4 | 8,194 | 8,049 |
| *Umbilicaria hispanica* | TBG_2322 | 38.71% | 130 | 3125324 | 96.8 | 43.4 | 9,111 | 9,021 |
| *Umbilicaria hispanica* | TBG_2337 | 54.22% | 60 | 4226768 | 97.3 | 41.9 | 8,781 | 8,696 |
| *Umbilicaria pustulata* | TBG_2333 | 33% | 26 | 2620629 | 97.3 | 37.7 | 9,569 | 9,503 |
| *Umbilicaria pustulata* | TBG_2345 | 32.26% | 31 | 2364512 | 96.8 | 35.7 | 8,790 | 8,740 |
| *Umbilicaria spodochoa* | TBG_2434 | 34.20% | 139 | 993216 | 97.0 | 40.9 | 8,791 | 8,705 |
| *Umbilicaria spodochoa* | TBG_2435 | 40.93% | 97 | 1249424 | 97.1 | 40.1 | 8,612 | 8,507 |
| *Umbilicaria subpolyphylla* | TBG_2323 | 41.14% | 190 | 1544375 | 99.6 | 58.2 | 16,993 | 16,852 |
| *Umbilicaria subpolyphylla* | TBG_2324 | 33.68% | 42 | 1514392 | 97.6 | 33.7 | 8,556 | 8,410 |
| *Umbilicaria grisea* | TBG_2336 | 42.54% | 40 | 1822796 | 96.9 | 44.43 | NA | NA |
| *Dermatocarpon miniatum* | TBG_2326 | 29.36% | 28 | 5077191 | 98.1 | 63.5 | 9,273 | 9,189 |
| *Dermatocarpon miniatum* | TBG_2331 | 26.28% | 22 | 4245366 | 98.4 | 49.8 | 7,938 | 7,871 |

**Figure 1.**
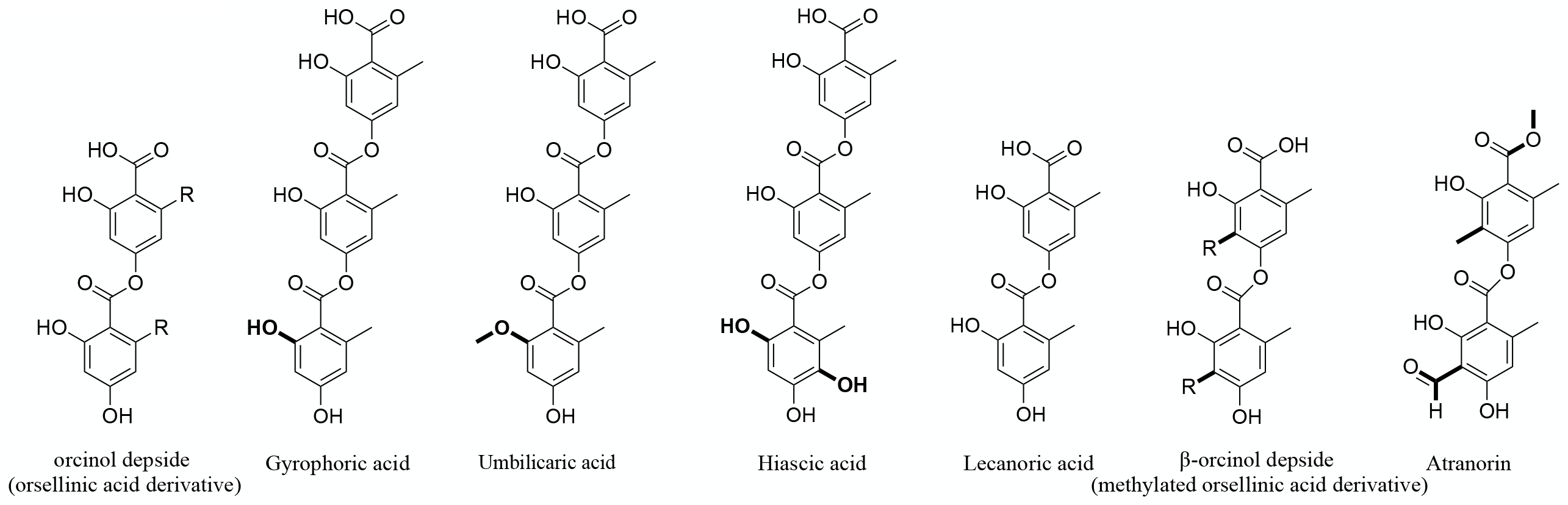
Chemical structures and nomenclature. Structure of a lichen depside, atranorin, GA and other depsides produced by *Umbilicaria* spp.

In the present study, we assembled highly contiguous long-read-based genomes of LFF of the genus *Umbilicaria*, identified the biosynthetic gene clusters of all the species and singled out candidate genes linked to GA biosynthesis.

## Materials and methods

### Sampling and dataset

We collected samples of the following eight *Umbilicaria* species: *U. deusta, U. freyi, U. grisea, U. subpolyphylla, U. hispanica, U. phaea, U. pustulata*, and *U. spodochroa* for genome sequencing (voucher information in Supplementary Table S1). When possible, we sequenced two samples of the same species collected in different climatic zones. This was done to take into account possible intra-specific variation in BGC content as recently shown in Singh et al. (34). The genome of *U. muhlenbergii* was downloaded from the JGI database. In addition, we sampled *Dermatocarpon miniatum* as a control, as it does not produce depsides/depsidones.

### DNA extraction, library preparation and genome sequencing

Lichen thalli were thoroughly washed with sterile water, and checked under the stereomicroscope for the presence of possible contamination and other lichen thalli. DNA was extracted from all the samples using a CTAB-based method (35) as presented in (36).

SMRTbell libraries were constructed according to the manufacturer’s instructions of the SMRTbell Express Prep Kit v. 2.0 following the Low DNA Input Protocol (Pacific Biosciences, Menlo Park, CA). Total input for samples was approximately 170-800 ng. Ligation with T-overhang SMRTbell adapters was performed at 20°C for 1 h or overnight. Following ligation, the libraries were purified with a 0,45 x or 0,8 x AMPure PB bead clean up step. The subsequent size selection step to remove SMRTbell templates <3 kb was performed with 2,2 x of a 40% (v/ v) AMPure PB bead working solution.

SMRT sequencing was performed on the Sequel System II with the Sequel II Sequencing Kit 2.0 using the continuous long read (CLR) mode or the circular consensus sequencing (CCS) mode, 30 h movie time with no pre-extension and Software SMRTLINK 8.0. Each metagenomic library was sequenced on one SMART cell at the Medical Center Nijmegen (the Netherlands), or at MPI Dresden.

### Genome assembly and annotation

The continuous long reads (i.e. CLR reads) from the PacBio Sequel II CLR run were first processed into highly accurate consensus sequences (i.e. HiFi reads) using PacBio tool CCS v5.0.0 with default parameters (https://ccs.how). HiFi reads were then assembled into contigs using the assembler metaFlye v2.7 (37). The resulting contigs were scaffolded with LRScaf v1.1.12 (github.com/shingocat/lrscaf, (38)). The scaffolds were then taxonomically binned to extract Ascomyocta reads with blastx using DIAMOND (--more-sensitive --frameshift 15 – range-culling) on a custom database and following the MEGAN6 Community Edition pipeline (39). All scaffolds assigned to Ascomycota were extracted as to represent the *Umbilicaria* spp. Assembly statistics such as number of contigs, total length and N50 were accessed with Assemblathon v2 (40) (Table 1). The completeness of the received mycobiont bins (i.e. the fungal genomes) was estimated using Benchmarking Universal Single-Copy Orthologs (BUSCO) analysis in BUSCO v4 (41).

### Identification and Annotations of Biosynthetic Gene Clusters

Functional annotation of genomes, including genes, proteins and BGC prediction (antiSMASH (antibiotics & SM Analysis Shell, v5.0)) was performed with scripts implemented in the funannotate pipeline (42, 43). First, the genomes were masked for repetitive elements, and then the gene prediction was performed using BUSCO2 to train Augustus and self-training GeneMark-ES (41, 44). Functional annotation was done with InterProScan (45), egg-NOG-mapper (46, 47) and BUSCO (41) with ascomycota_db models. Secreted proteins were predicted using SignalP (48) as implemented in the funannotate ‘annotate’ command.

### Selecting candidate gene clusters linked to GA biosynthesis

We used the following criteria to select the candidate gene cluster associated with GA synthesis in *Umbilicaria*: 1) it must contain a NR-PKS (as some of the structural features of a NP can be directly inferred from the domain architecture of the *PKS*: *PKSs* without reducing domains (*NR-PKSs*) are linked to non-reduced compounds such as gyrophoric acid, olivetoric acid (16), physodic acid (16) and grayanic acid (15)), 2) it must be present in all the *Umbilicaria* genomes, as all the species have GA as the major secondary metabolite (33), and 3) it must be closely related to the *PKSs* involved in the synthesis of orsellinic acid-based compounds (15, 16), because orsellinic acid units constitute the building blocks of GA.

### Phylogenetic analyses

We extracted the amino acid sequences of all the NR-PKS from the BGCs predicted by the antiSMASH for all the *Umbilicaria* species and *Dermatocarpon miniatum* (Supplementary Table S2). Additionally, the NR-PKS sequences of the following species were downloaded from the previous publications and public databases: *Cladonia borealis, C. grayi, C. macilenta, C. metacorallifera, C. rangiferina, C. uncialis, Pseudevernia furfuracea, Stereocaulon alpinum* and *Umbilicaria muhlenbergii*. The final dataset contains amino acid sequences of 229 NR-PKSs from 18 species belonging to four LFF genera. The sequences were aligned using MAFFT as implemented in Geneious v5.4 (49, 50). Gaps were treated as missing data. The maximum likelihood search was performed on the aligned sequences with RAxML-HPC BlackBox v8.1.11 (51) on the Cipres Scientific gateway (52). Phylogenetic trees were visualized using iTOL (53).

### BGC clustering and novel BGCs: BiG-SCAPE and CORASON

We used BiG-SCAPE and CORASON (54) to identify the gene cluster networks and infer evolutionary relationships among clusters of interest among different *Umbilicaria* spp. BiG-SCAPE utilizes antiSMASH (42) and MIBiG databases (55) for inferring BGC sequence similarity networks, whereas CORASON employs a phylogenomic approach to infer evolutionary relationships between the clusters. BiG-SCAPE v1.0.1 was run in --auto mode, to identify BGC families using antiSMASH output files (.gbk) as input. Networks were generated using similarity thresholds of 0.25. The most likely GA cluster from all the *Umbilicaria* spp. was examined for conservation and variation among different *Umbilicaria* species using CORASON pipeline. The antiSMASH .gbk files of the corresponding clusters, based on phylogenetic grouping, were used as input. The most-likely GA cluster from *U. deusta* was used as reference to fish out the most closely-related clusters from the other *Umbilicaria* spp.

## Results

### Genome sequencing, assembly and annotation

The genome quality stats and assembly reports of all the genomes generated for this study are presented in Table 1.

### Phylogenetic analysis

To search for PKS genes involved in the synthesis of GA, we performed a phylogenetic analysis by incorporating our sequences into the most comprehensive PKS dataset currently available (Supplementary Table S2) (13, 16). NR-PKSs have been categorized into nine groups based on protein sequence similarity and PKS domain architecture (13). We identified a total of 110 NR-PKSs that were present in 15 *Umbilicaria* genomes (12 NR-PKSs on average per species). Four NR-PKSs were common to all species: PKS15, PKS16, PKS20 and a novel PKS clade (forming a monophyletic, supported clade to PKS33, Fig. 2). Only one NR-PKS per species formed a supported monophyletic clade with PKS16 (Group I, i.e., orsellinic acid, depside and depsidone NR-PKSs) (Fig. 2). The most-likely NR-PKS for the depsidone grayanic acid and the depsides olivetoric and physodic acid fall within this PKS clade.

**Figure 2.**
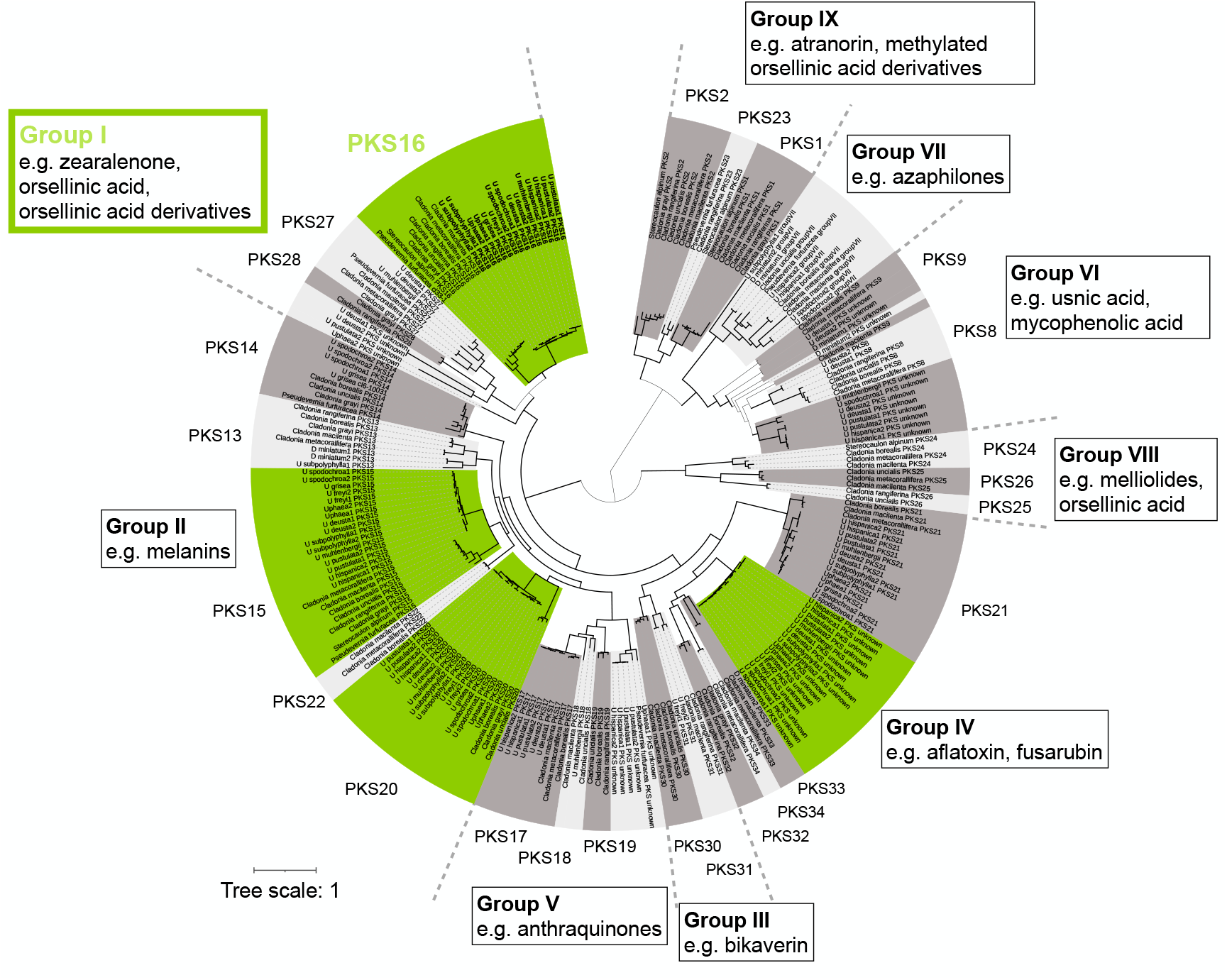
NR-PKS phylogeny of lichen-forming fungi. This is a maximum-likelihood tree based on amino acid sequences of NR-PKSs from nine *Umbilicaria* spp., six *Cladonia* spp., *Dermatocarpon miniatum, Stereocaulon alpinum* and *Pseudevernia furfuracea*. Branches in bold indicate bootstrap support >70%. Green color clades represent the PKSs common to all nine *Umbilicaria* spp. used in this study. PKS groups are based on Kim et al. (13).

### Gyrophoric acid cluster

The cluster most likely associated with GA is the cluster containing PKS16 (Fig. 2), as 1) it is present in all *Umbilicaria* spp., 2) it contains an *NR-PKS* and 3) it forms a monophyletic group with the clade “Group I, PKS 16” from Kim et al (11). This cluster contains about 11-15 genes, including the core biosynthetic gene *NR-PKS* and a c*yt P450* (Fig. 3). The other genes code for unidentified proteins. The *U. deusta* PKS16 cluster is presented as an example of GA cluster (Fig. 3). The *PKS* is present in all *Umbilicaria* species investigated and displays high homology across species.

**Figure 3.**
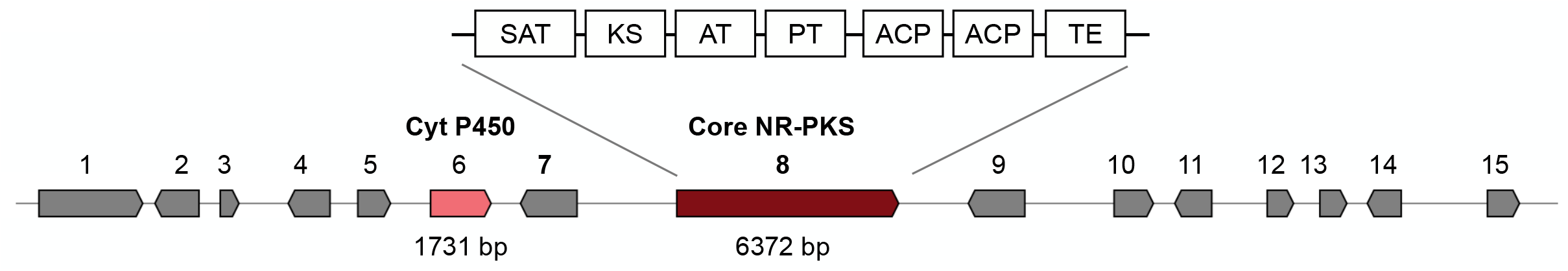
Gyrophoric acid cluster from *Umbilicaria deusta* as predicted by antiSMASH. Colored boxes indicate genes. Genes in grey represent genes coding for unknown proteins

### BGC clustering: BiG-SCAPE and CORASON

BGC sequence similarity networks group gene clusters at multiple hierarchical levels. This analysis implements a ‘glocal’ alignment mode that accurately groups both complete and fragmented BGCs. The BGCs forming a supported monophyletic clade to PKS16 (Group I) were then analyzed for conservation across species using CORASON. The CORASON analysis showed that only three genes on the cluster were shared among the studied *Umbilicaria* species: the core *PKS* and the two genes of unknown function/proteins adjacent to the core gene (Fig. 4).

**Figure 4.**
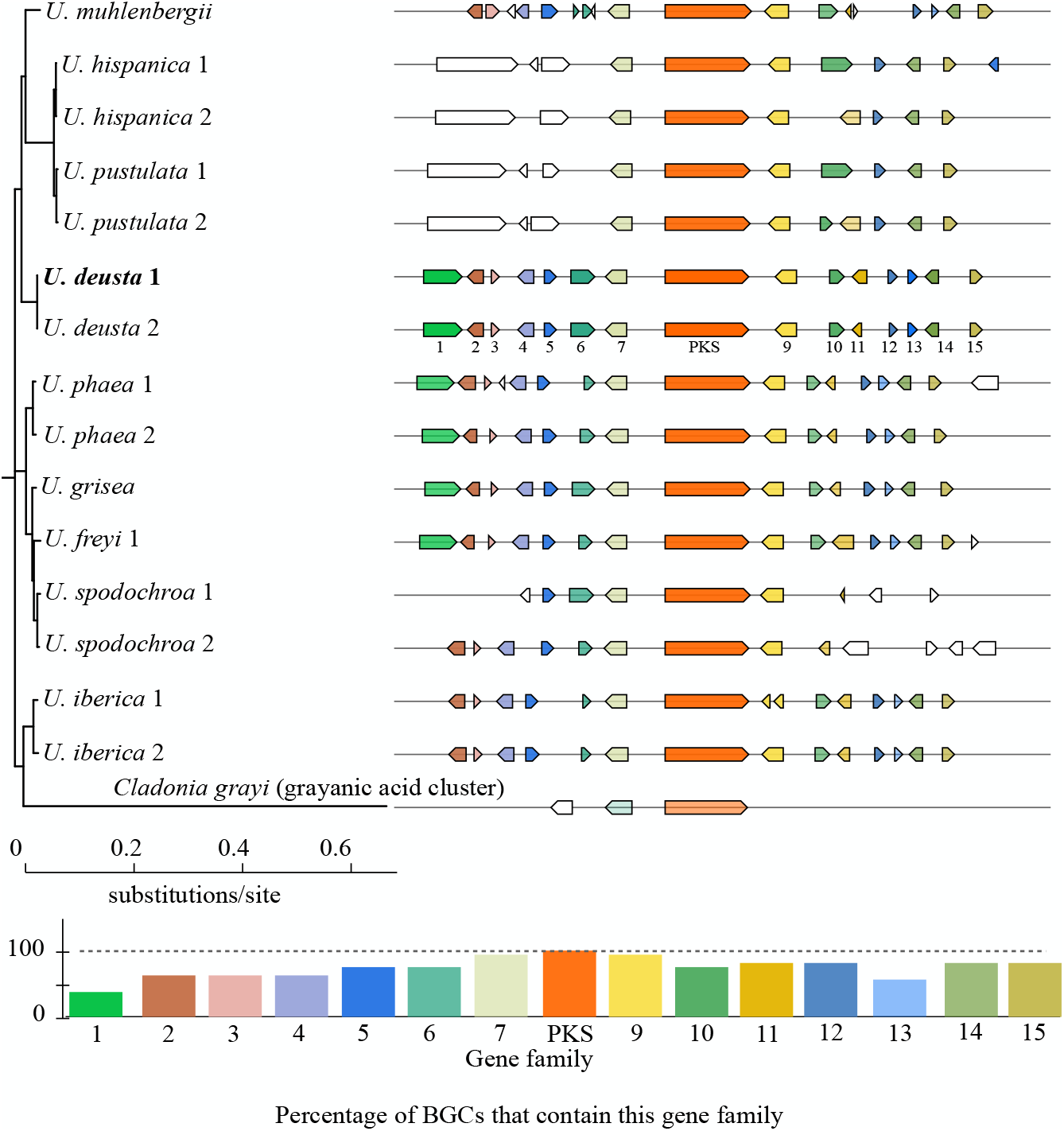
CORASON-based PKS phylogeny to elucidate evolutionary relationships and cluster organization of GA cluster in *Umbilicaria* spp. The bar plot below depicts the percentage of *Umbilicaria* species in which a particular gene is present.

## Discussion

### Gyrophoric acid PKS

We found only one PKS per species forming a supported monophyletic clade to PKS16 (Group I, i.e., orsellinic acid and depside/depsidone PKSs) (Fig. 2). These are the most likely GA PKSs.

The BGC associated with the biosynthesis of the following lichen depsides and depsidones have been identified so far: atranorin (13), lecanoric-(14), grayanic-(15), olivetoric-(16) and physodic acid (16). All these studies demonstrate that the PKS alone is capable of synthesizing the backbone depside, whereas modifications such as methylation and oxidation are made by enzymes coded by other genes in the cluster after the release of the depside from the PKS. For instance, the synthesis of atranorin involves at least three genes present within the atranorin cluster, but the core depside is coded only by the PKS (13). The other two genes, a carboxyl methylase (O-methyltransferase) and a *cyt P450*, methylate the carboxyl group and oxidize the methyl group (into -CHO), respectively, to produce the final product atranorin (Fig. 1). As GA does not have side chain modifications (Fig. 1) we propose that the *PKS* alone is involved in GA synthesis.

The depside PKSs identified so far code for didepsides, i.e. they contain two phenolic rings joined with an ester bond, for example gyrayanic acid, atranorin, physodic acid and olivetoric acid (13, 15, 16). Ours is the first study to identify the most-likely PKS associated with a tridepside synthesis, i.e., three phenolic rings joined with two ester bonds. Our study suggests that the *PKSs* coding for a didepside and a tridepside differ only in the length of the sequence of the SAT domain. A tridepside *PKS* contains longer SAT coding sequence than the didepside *PKS*. The number of ACP and PT domains is the same between the two.

### GA cluster in *Umbilicaria* spp

The most-likely GA cluster contains about 11-15 genes in different *Umbilicaria* spp. (Fig. 3, 4). Interestingly, only three genes are common to all analyzed species, the *PKS* and two genes of unknown function (with low sequence similarity to known genes) upstream and downstream of the *PKS I* (Fig. 4). This suggests that these three genes form an integral part of the GA cluster, whereas the other genes are facultative among GA producers. Differences among the clusters synthesizing the same compound have been reported before, and have been associated with species-specific BGC regulation or modifications to the depside released by the PKS (26, 56).

The most-likely GA cluster also contains a *cyt P450* (Fig. 3, 4), which has been associated with depsidone production or oxidation of an acyl chain (13, 15, 16). However, the location and orientation of the *cyt P450* in the putative *Umbilicaria* GA cluster is different from a typical depsidone cluster (Fig. 3) (15). In the GA cluster, the *cyt P450* is not located next to and has the same orientation as the *PKS*, whereas in a depsidone cluster, *cyt P450* lies next to the *PKS*, in opposite orientation. Such organization is suggestive of genes being regulated and co-expressed by the same promoter (15). This is the case for the depsidone grayanic acid synthesis (in *Cladonia grayi*), which involves the synthesis of the depside intermediate by *PKS* followed by oxidation of the released depside into depsidone (15). The *PKS* and *cyt P450* form the integral part of depsidone synthesis (57) whereas the depside is coded by the *PKS* alone, with the exception of the side chain modifications (13, 14). Therefore, despite being part of the GA cluster, the *cyt P450* does not seem to be involved in GA synthesis or in the synthesis of umbilicaric- and/or lecanoric acid reported from *Umbilicaria* spp. analyzed in this study. The synthesis of hiascic acid however would require the hydroxylation of a methyl group by cyt P450 enzyme after the depside is released from the PKS (Fig. 1, the OH group in bold in hiascic acid). The lower proportion of hiascic acid as compared to GA could be because the *cyt P450* may not be co-expressed with the *PKS*.

Even though the functions of most of the genes identified in the present study are unknown, our study provides novel insights into GA cluster composition and organization across different species (Fig. 4). This information is crucial in order to open the way for future genetic manipulation of the GA biosynthetic pathway that may be aimed at increasing structural diversity and/or yield of the products, as well as in order to generate analogs with novel properties.

### One cluster, different compounds

Variation in cluster composition reflects the potential to produce diverse NPs. Apart from GA, other depsides related in structure to GA, i.e., lecanoric-, umbilicaric- and hiascic acid (Fig. 1) are often reported in *Umbilicaria* spp. as minor metabolites (31). Interestingly, we found only one orcinol-depside *PKS* in *Umbilicaria* spp (Fig. 2). This strongly indicates that all the *Umbilicaria* depsides are coded by the same PKS cluster. One cluster coding for different, structurally-related compounds has also been reported previously (16, 56, 58). For instance, in the case of the antifungal drug caspofungin acetate, a semisynthetic derivative of the NP pneumocandins from the fungus *Glarea lozoyensis*, selective inactivation of different genes in this biosynthetic gene cluster generates 13 different analogues, some of them with elevated antifungal activity relative to the original compound and its semisynthetic derivative (59). Similarly, the aspyridone biosynthetic cluster from *Aspergillus nidulans* produces eight different compounds in a heterologous host (58). These studies show that a single PKS cluster is capable of producing different compounds depending upon which genes are co-expressed and on the available starters. In lichens, a single *PKS* has been associated with the synthesis of olivetoric and physodic acid (16) and the same *PKS* has been shown to be involved in the synthesis of lecanoric acid in a heterologous host (14). We propose that the same PKS cluster is most likely involved in the synthesis of GA, umbilicaric- (an additional methyl group, Fig. 1), hiascic- (additional hydroxyl group, Fig. 1), and lecanoric acid (didepside with no side chains, Fig. 1) in *Umbilicaria*. It is possible, however, that in nature only GA is synthesized in members of the genus *Umbilicaria*, and the co-occurring minor compound lecanoric acid is a hydrolysis product of GA (57).

Interestingly, although umbilicaric acid is reported from some *Umbilicaria* species (*U. grisea, U. freyi, U. mühlenbergii* and *U. subpolyphylla* (31, 61)), O-methyltransferase (OMT) was not identified in the depside-related BGC of any *Umbilicaria* species (Fig. 3). OMT would be required for the methylation of oxygen to produce umbilicaric acid (Fig. 1). Its absence from depside-related BGCs suggests that an external OMT, e.g. from other BGCs, might be involved in the production of umbilicaric acid in *Umbilicaria*. This could explain the lower amounts of umbilicaric acid as compared to GA found in these species (31). In contrast, when the O-methylated compound is the major secondary metabolite, as in the case of grayanic acid and atranorin, OMT is an integral part of the BGC and is co-expressed along with the other crucial genes for grayanic acid production, i.e., the *PKS* and *cyt P450* (13, 15).

### Future perspectives

Advances in long-read sequencing and in computational approaches to genome mining not only enable linking biosynthetic genes to NPs but also provide an overview of the entire gene cluster composition and organization. Ours is the first study to identify the most-likely GA cluster, which is essential for opening up avenues for biotechnological approaches to producing and modifying this compound and possibly other lichen compounds. In particular, this information can be applied to generate novel NP analogs with improved pharmacological properties via synthetic biology, biotechnology and combinatorial biosynthesis approaches. This paves the way to an entirely new horizon for utilizing these understudied taxa for pharmacological industry and drug discovery.

## Acknowledgements

We thank Prof. Daniele Armaleo (Duke University) for his inputs on the domain composition of didepsides and tridepsides.

## Supplementary Tables

1. Voucher information table of the *Umbilicaria* samples collected for this study.
2. Dataset used for the phylogenetic analysis, along with the amino acid sequences of the PKS

